# β-catenin Signaling Regulates Cell Fate Decisions at the Transition Zone of the Chondro-Osseous Junction During Fracture Healing

**DOI:** 10.1101/2020.03.11.986141

**Authors:** Sarah Anne Wong, Diane Hu, Tiffany Shao, Erene Niemi, Emilie Barruet, Blanca M Morales, Omid Boozarpour, Theodore Miclau, Edward C Hsiao, Mary Nakamura, Chelsea S Bahney, Ralph S Marcucio

## Abstract

Chondrocytes within the fracture callus transform into osteoblasts during bone regeneration, but the molecular mechanisms regulating this process are unknown. Wnt ligands are expressed within the fracture callus, and hypertrophic chondrocytes undergoing transformation to osteoblasts exhibit nuclear localization of β-catenin, indicating active Wnt signaling in these cells. Here, we show that conditional knock out (cKO) of β-catenin in chondrocytes inhibits the transformation of chondrocytes to osteoblasts, while stabilization of β-catenin in chondrocytes accelerates this process. After cKO, chondrocyte-derived cells were located in the bone marrow cavity and upon re-fracture formed cartilage. Lineage tracing in wild type mice revealed that in addition to osteoblasts, chondrocytes give rise to stem cells that contribute to repair of subsequent fractures. These data indicate that Wnt signaling directs cell fate choices of chondrocytes during fracture healing by stimulating transformation of chondrocytes to osteoblasts, and provide a framework for developing Wnt-therapies to stimulate repair.

## Introduction

Fractures heal through the process of endochondral ossification, in which cartilage forms between the fractured bone ends and transforms into bone.^1–3^ This process begins with differentiation of progenitor cells derived from the periosteum into chondrocytes, which proliferate to form the cartilage callus.^4–6^ As chondrocytes mature to a hypertrophic state, they mineralize their matrix and induce vascular invasion.^7–11^ Genetic lineage tracing studies demonstrate that hypertrophic chondrocytes directly contribute to the formation of bone by transforming into osteoblasts.^8,12–17^ This transformation has been shown using a variety of genetic models during bone development, postnatal growth, and repair in the appendicular and craniofacial skeleton.^8,12–17^ However, the molecular signals that regulate this process during bone healing are largely unknown.^8,16^

Evidence demonstrates that the canonical Wnt pathway may play a key role in regulating chondrocyte-to-osteoblast transformation during endochondral repair. Loss- and gain-of-function experiments modifying Wnt signaling have demonstrated the necessity of this pathway for chondrocyte transformation during long-bone development.^16^ In the context of endochondral repair, hypertrophic chondrocytes located at the chondro-osseous border of the fracture callus, where chondrocyte-to-osteoblast transformation occurs, have been shown to have nuclear localization of β-catenin, indicating these cells undergo active Wnt signaling.^8^

The canonical Wnt pathway requires the transcriptional co-activator β-catenin to mediate its effect in modulating gene expression.^3,18^ β-catenin is constitutively expressed. However, when the pathway is inactive, β-catenin is targeted for ubiquitin-mediated degradation by the “destruction complex.” When the canonical Wnt pathway is activated the destruction complex is disrupted, stabilizing β-catenin in the cytoplasm and allowing it to translocate to the nucleus where it binds to members of the T-cell factor/lymphocyte elongation factor (TCF/LEF) family to regulate Wnt target gene expression.^3,18^

Due to the Wnt pathway’s known role in promoting direct bone formation numerous Wnt-based therapies are currently being developed.^1^ The hydrophobic nature of Wnt ligands and requirement of palmitoylation for biological activity precludes treatment with recombinant Wnts as a direct clinical therapeutic.^1,19–22^ Thus, the majority of Wnt-based therapies aim to activate Wnt signaling by neutralizing pathway inhibitors.^1,23^ One promising Wnt pathway regulator, Romosozumab, is a monoclonal antibody that works by binding and inhibiting the activity of sclerostin.^1,23,24^ Understanding endogenous Wnt signaling during endochondral fracture repair is critical to effectively optimize the timing and targets of Wnt therapeutics for bone regeneration. In this paper we demonstrate that activation of canonical Wnt signaling during chondrocyte-to-osteoblast transformation is necessary for normal fracture repair. This evidence provides key insight into the process of endochondral repair and advances the development of novel Wnt-based fracture therapies.

## Results

### Inhibition of Canonical Wnt Signaling in Fracture Callus Chondrocytes Significantly Reduces Bone Formation During Repair

To determine the role of canonical Wnt signaling in regulating chondrocyte-to-osteoblast transformation, the pathway was conditionally inhibited in chondrocytes by inducing deletion of β-catenin using the Aggrecan-Cre^ERT2^ mouse. Cre recombination was induced with daily injections of Tamoxifen from 6 to 10 days post-fracture, which has been previously shown to induce robust Cre recombination in chondrocytes.^8,13^ Cartilage formation at early stages of repair was unaffected (D7 & D10). However, significantly reduced bone formation and increased cartilage retention occurred at later time points (D14 & D21) during which chondrocyte transformation primarily occurs **(Fig 1, S1)**. A statistically significant reduction in bone formation was observed with deletion of either one or both copies of β-catenin. However, the reduction in bone formation was greater with the loss of both copies. **(Fig S2)**.

**Figure 1:**
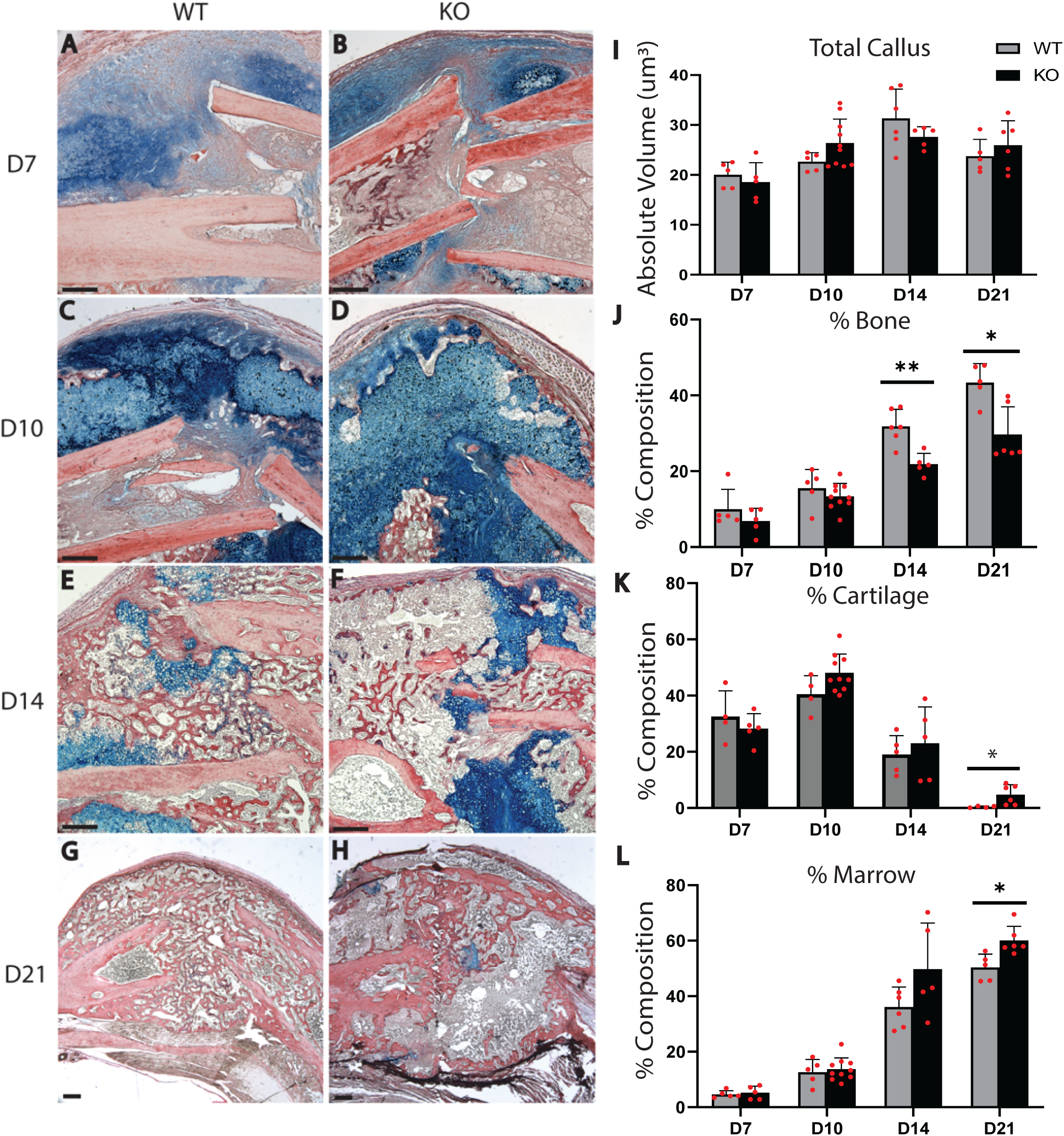
Deletion of β-catenin in Chondrocytes Significantly Impairs Endochondral Healing. **(A-H)** HBQ histology and **(I-L)** stereological quantification of fracture callus size and composition shows that inhibition of canonical Wnt signaling through conditional deletion of β-catenin in chondrocytes significantly reduces trabecular bone formation **(E-H, J)** and increases cartilage retention **(G-H, K)** at later time points (D14, D21). Callus size and composition are unaffected at early time points (D7, D10). Stereological quantification of Callus size and composition are unaffected at early time points (D7, D10). HBQ histology: cartilage = blue, bone = red. N=5/group/time point. Scale bar = 1000μm. (*) = p< 0.05. (**) = p< 0.01.

The above experiments were conducted using male mice. To confirm that the canonical Wnt pathway plays a similar role in female mice, females were harvested at 14 days post-fracture. Analysis revealed that deletion of β-catenin in chondrocytes of female mice produced a similar reduction in bone formation and increase in cartilage retention **(Fig S3, A)** as seen in male mice **(Fig 1, E-F, J-K)**. Furthermore, comparison of male and female mice revealed that although females had significantly lower body weight than males **(Fig S3, B)**, there was no statistically significant difference between sexes for tissue composition parameters when normalized to body weight **(Fig S3, C-D)**, except for a slight increase in total callus size in females when β-catenin was deleted **(Fig S3,C)**.

TUNEL staining revealed that deletion of β-catenin did not significantly increase chondrocyte cell death. Indeed, lineage tracing analysis using a tdTomato reporter showed survival of chondrocytes as detached cells filling the marrow space **(Fig 2)**. tdTomato-positive cells were observed within the marrow cavity as late as 35 days post-fracture **(Fig 2, J, N)**. The specificity of tdTomato fluorescence was confirmed by the complete absence of expression in marrow cells found at the growth plate of reporter mice **(Fig 2, E, H)** and by the lack of tdTomato expression in all cells comprising the fracture calli of mice lacking the tdTomato reporter **(Fig 2, F, I)**.

**Figure 2:**
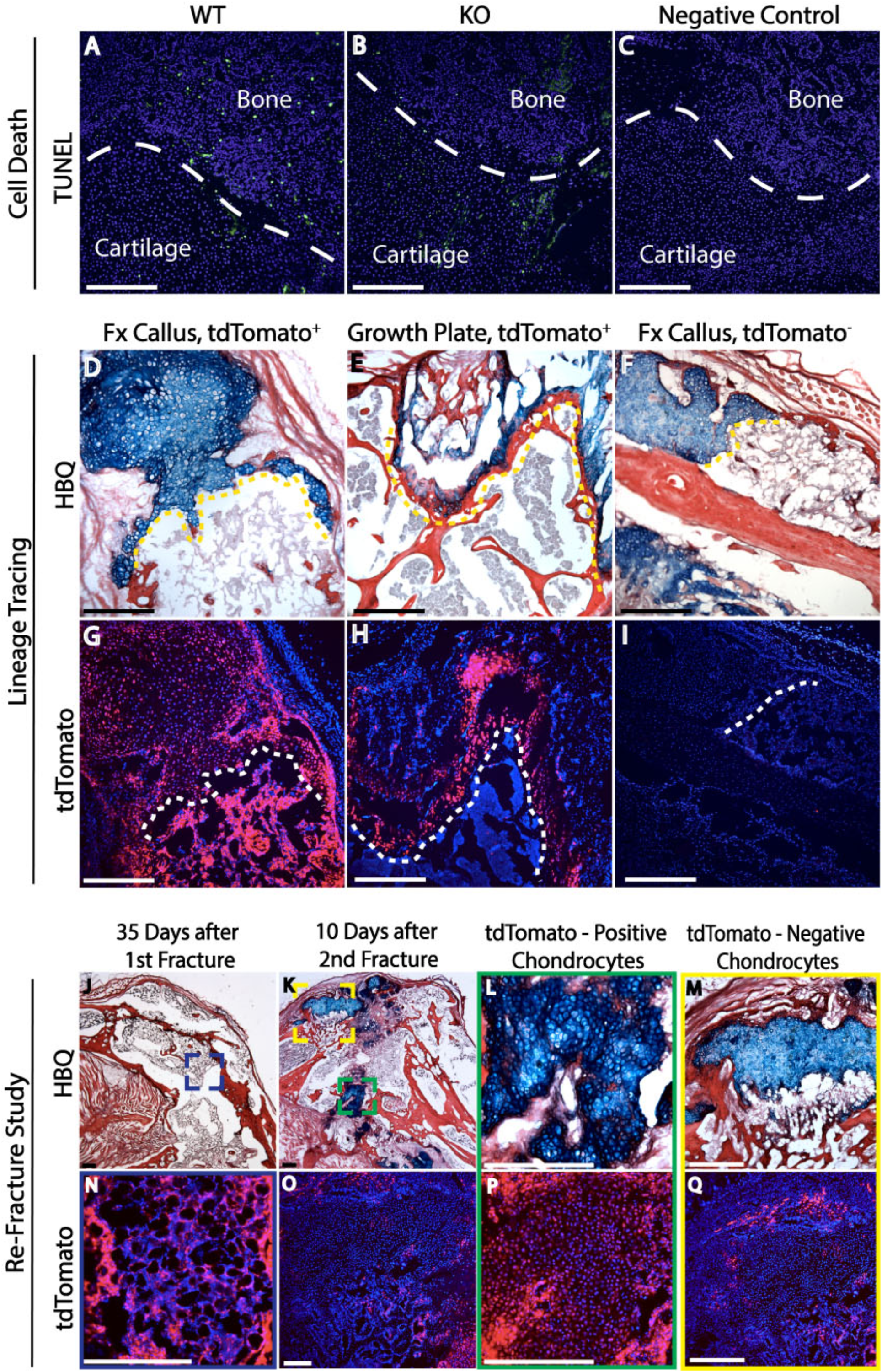
Chondrocytes Survive within the Marrow Cavity Following Deletion of β-catenin. **(A-C)** TUNEL analysis at 14 days post-fracture demonstrates that inhibition of canonical Wnt signaling in chondrocytes through deletion of β-catenin does not significantly increase chondrocyte cell death. **(D-I)** Furthermore, lineage tracing analysis using a tdTomato reporter at 14 days post-fracture reveals that chondrocytes survive within the marrow cavity following deletion of β-catenin. **(E, H)** The specificity of tdTomato fluorescence was confirmed by its complete absence in marrow cavity cells found at the growth plate of reporter mice as well as **(F, I)** in all cells of fracture calli from wildtype mice lacking the tdTomato reporter. **(J)** No cartilage was observed 35 days post-fracture. **(N)** However, the marrow cavity contained a robust population of tdTomato-positive cells. In another set of mice, a second fracture was created in the same anatomical location 35 days following initial fracture and samples were harvested 10 days later. (**K-M)** HBQ histology reveals several small islands of cartilage forming a new cartilage callus. **(O-Q)** tdTomato lineage tracing reveals that some of the cartilage islands contain **(L, P)** tdTomato-positibe cells whereas **(M, Q)** others are tdTomato-negative. HBQ histology: cartilage = blue, bone = red. DAPI counterstain was used to visualize nuclei (blue). N=5. Scale = 200μm.

Loss of β-catenin resulted in unique changes in cellular morphology at the chondro-osseous border (“Transition Zone”). Unlike controls, callus cartilage did not directly connect to trabecular bone. Rather, callus chondrocytes appeared to “slough off” into the marrow space, a process potentially aided by TRAP-positive cells, which exhibited increased localization to the borders of the cartilage callus **(Fig S4)**. Interestingly, a population of FACS-analyzed tdTomato+ marrow cells was positive for CD45, a marker of hematopoietic cell lineages **(Fig 3, B)**.^25,26^ In contrast, marrow cells from control mice expressing the tdTomato reporter but possessing wildtype copies of β-catenin demonstrated neither significant tdTomato fluorescence nor significant tdTomato, CD45 dual-labelling **(Fig 3, C)**. CD45 staining did not bleed into the tomato channel as evidenced by minimal CD45 staining observed in unstained cells **(Fig 3, A, E).** Furthermore, the degree of tdTomato, CD45 dual-labelling in control mice **(Fig 3, C;** β-catenin+/+, tdTomato+) was comparable to that seen in wildtype mice **(Fig 3, D;** β-catenin+/+, tdTomato-), which did not possess the tdTomato reporter. Together this data indicate that callus chondrocytes do not, under normal conditions, remain detached within the marrow cavity and that deletion of β-catenin in chondrocytes promotes hematopoietic engulfment of some cells.

**Figure 3:**
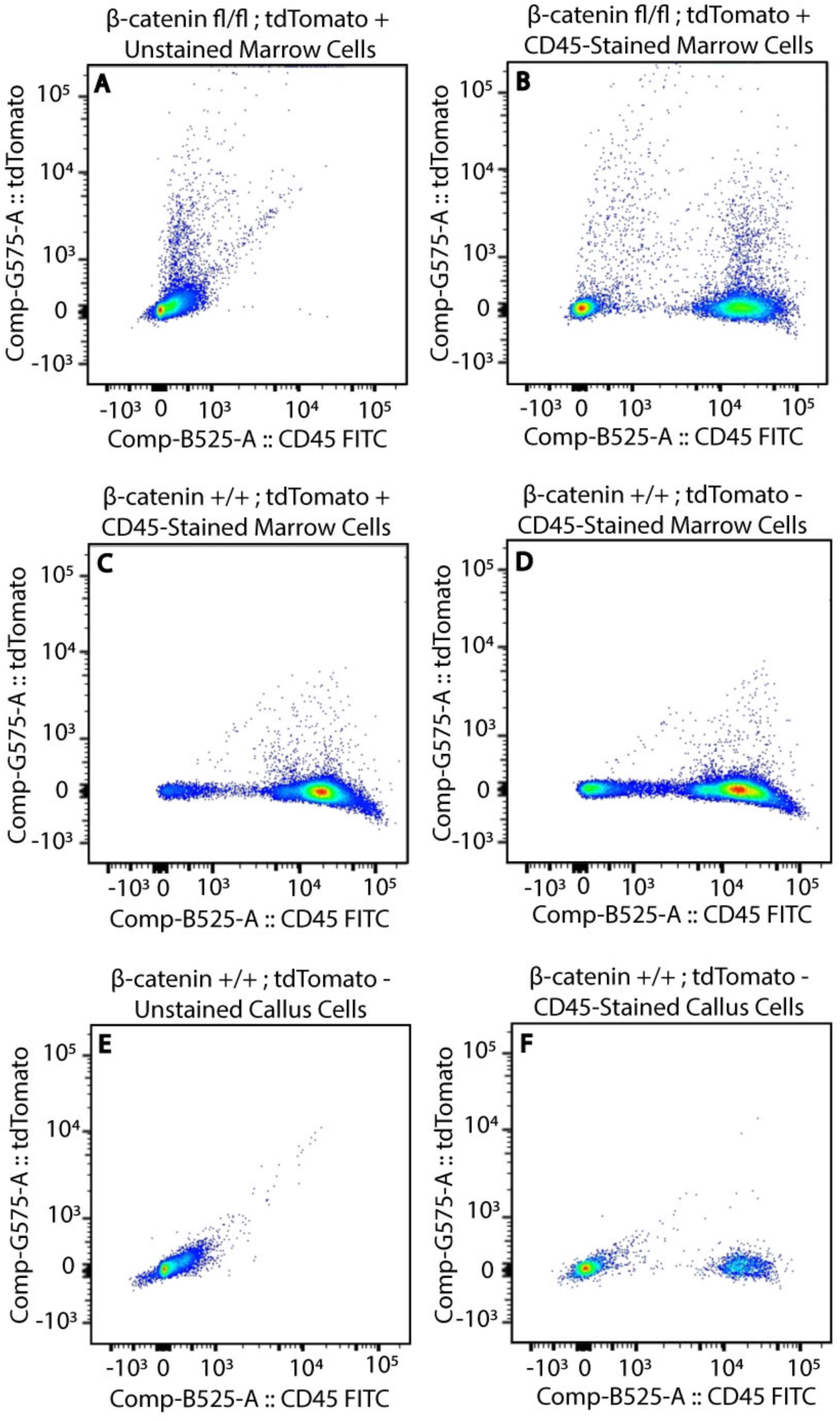
FACS Analysis of tdTomato+ Cells Reveals Chondrocyte Engulfment by Phagocytic Hematopoietic Cells. **(B)** FACS analysis reveals that deletion of β-catenin results in a significant population of tdTomato+ marrow cells that are also positive for CD45. **(C)** In contrast, FACS-analysis of marrow cells from control mice (β-catenin +/+, tdTomato+) demonstrates neither significant tdTomato+ fluorescence nor significant tdTomato+, CD45+ dual-labelling. **(A, E)** CD45 staining did not bleed into the tomato channel as evidenced by minimal CD45 staining observed in unstained cells**. (C)** Furthermore, the degree of tdTomato, CD45 dual-labelling in control mice (β-catenin+/+, tdTomato+) was comparable to that seen in **(D)** wildtype mice (β-catenin+/+, tdTomato-). Together this data indicate that callus chondrocytes do not, under normal conditions, remain detached within the marrow cavity and that deletion of β-catenin in chondrocytes promotes their engulfment by hematopoietic cells. N=3/cell population.

However, not all tdTomato-positive cells were engulfed. To test the extent to which skeletogenic potential was retained in these cells, we created new fractures in these mice over the site of the original injury. By day 35 the callus was completely devoid of cartilage and the marrow cavity contained a robust population of detached tdTomato-positive cells that appeared mesenchymal and not embedded in a matrix **(Fig 2, J, N)**. Upon refracture at 35 days post-fracture and harvest 10 days later, tdTomato-positive cells were observed as chondrocytes embedded in newly formed cartilage matrix in the fracture calli, indicating that deletion of β-catenin does not inhibit differentiation along the chondrocyte lineage. The specificity of the reporter in labelling chondrocytes that had previously undergone β-catenin deletion was confirmed in that not all chondrocytes within the re-fracture calli were tdTomato-positive. **(Fig 2, K-M, O-Q)**.

### Over-activation of Canonical Wnt Signaling in Fracture Callus Chondrocytes Accelerates Bone Repair by Promoting Chondrocyte Transformation

The converse, gain-of-function experiment was performed in which an indestructible, stabilized form of β-catenin was induced to be conditionally expressed in chondrocytes starting six days post-fracture. RT-PCR analysis of Wnt target gene *Axin2* expression in fracture calli harvested 10 days post-fracture confirmed that canonical Wnt signaling was over-activated **(Fig 4, M)**. Wnt pathway gain-of-function not only increased the absolute volume and percent callus composition of bone compared to controls, but it also accelerated bone formation as evidenced by the shift in bone formation to earlier time points (D10) **(Fig 4, O)**. This increase in bone formation corresponded with a simultaneous significant reduction in callus cartilage composition **(Fig 4, P)**. Together, these data suggest that stimulation of canonical Wnt signaling in chondrocytes results in accelerated chondrocyte-to-osteoblast conversion **(Fig 4 & S5)**.

**Figure 4:**
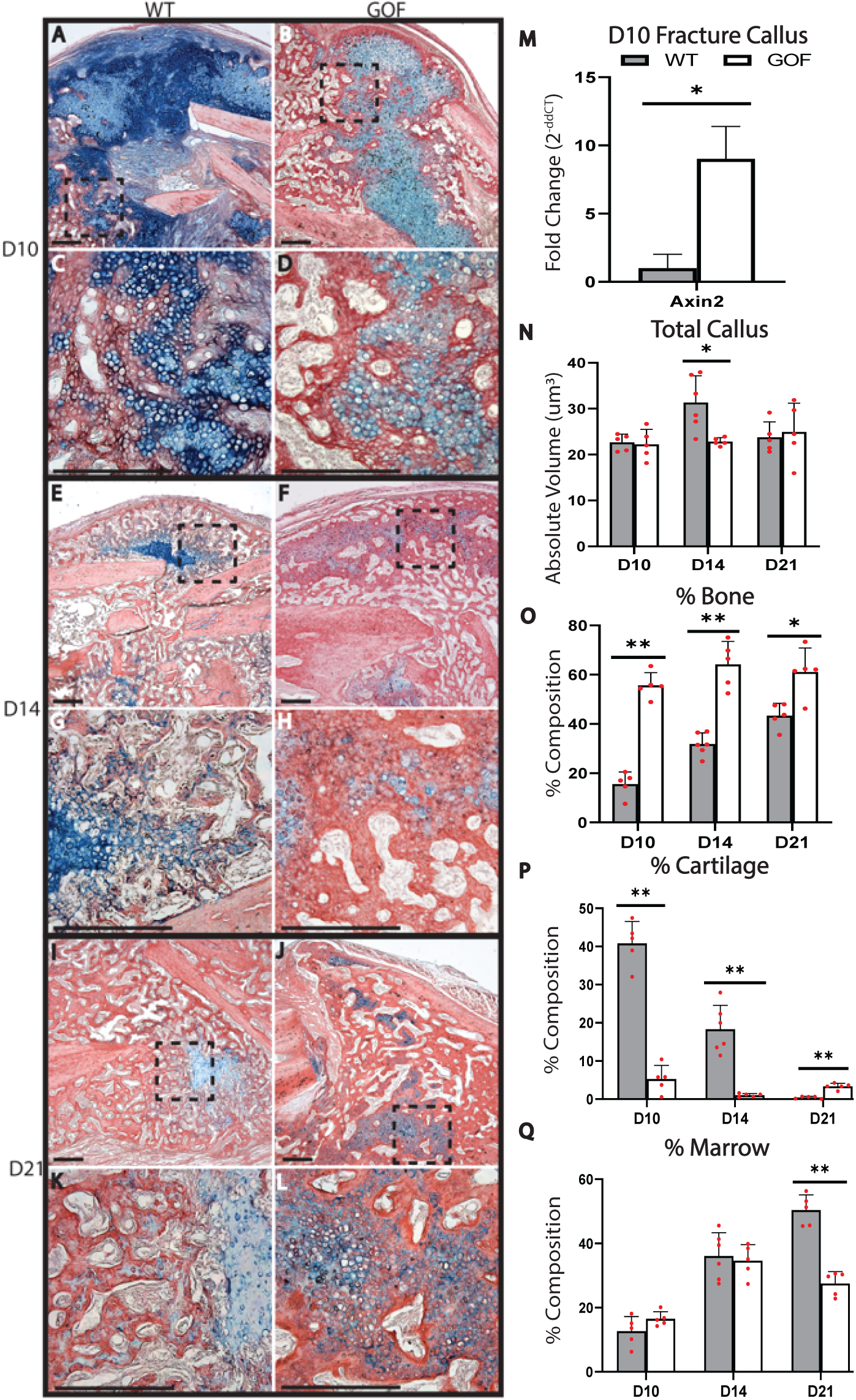
Overactivation of Canonical Wnt Signaling through Induced Stabilization of β-catenin in Chondrocytes Significantly Increases Bone Formation and Accelerates Endochondral Healing. **(M-Q)** Stereological quantification of **(A-L)** HBQ histology reveals that over-activation of canonical Wnt signaling through the conditional stabilization of β-catenin in chondrocytes **(N)** significantly increases and accelerates trabecular bone formation at early time points (D10). **(P)** This increase in bone formation occurs simultaneously with a significant reduction in callus cartilage composition. HBQ histology: cartilage = blue, bone = red. N=5/group/time point. Scale = 1000μm. (*) = p< 0.05. (**) = p< 0.01.

### Macrophages, Chondrocytes, and Endothelial Cells Express Wnts within the Fracture Callus

The Transition Zone at the chondro-osseous boarder of the fracture callus is hallmarked by the invasion of blood vessels, **(Fig 5, A)**, and we have previously shown that paracrine signaling from endothelial cells can stimulate chondrocytes to mineralize their surrounding matrix.^1,8,14^ In order to understand whether this may be due to endothelial cell secretion of Wnts, we treated 293-STF luciferase Wnt reporter cells with 20% human umbilical vein endothelial cell (HUVEC)-conditioned media. HUVEC-conditioned media produced the same level of Wnt pathway activation as treatment with 2nM Wnt3a and significantly higher activation compared to mock treatment **(Fig 5, B)**. Furthermore, RT-PCR analysis of HUVECs demonstrated that these cells express Wnts 3 and 7b as well as R-spondin 3, one of the four R-spondin secreted agonists responsible for the potentiation of Wnt signaling **(Fig 5, C)**.^27^ Thus, endothelial cells are not only capable of releasing Wnts but are also able to activate canonical Wnt signaling.

**Figure 5:**
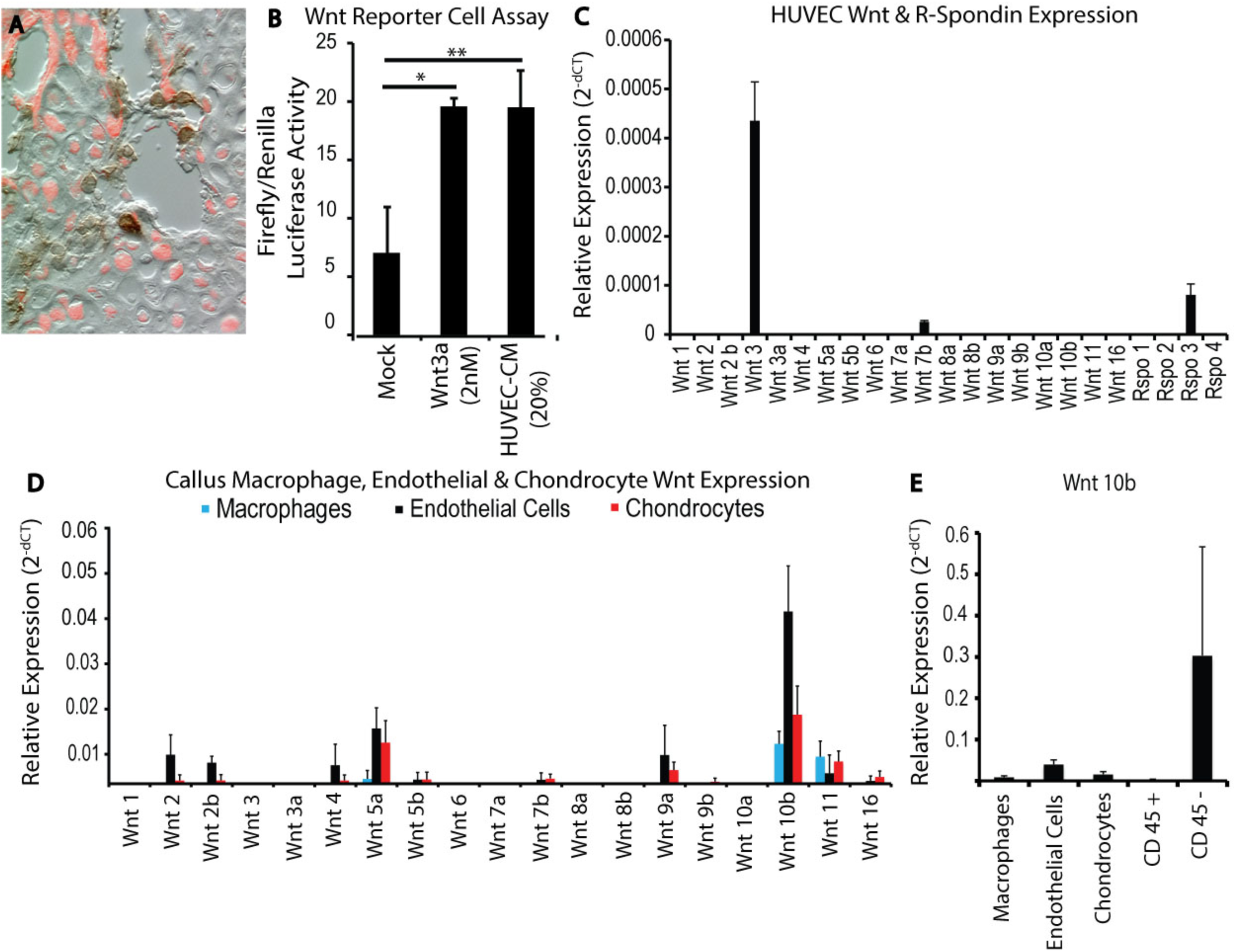
In Vitro and In Vivo Wnt Pathway Analysis. **(A)** The Transition Zone is hallmarked not only by the chondro-osseous border where callus cartilage joins the newly-formed bone but also by the presence of macrophages and invading blood vessels (tdTomato=chondrocytes, brown=F4/80 IHC). **(B)** Secreted factors from human umbilical vein endothelial cells (HUVECs) were capable of activating canonical Wnt signaling as evidenced by the significant increase in luciferase activity of 293-STF Wnt reporter cells treated with 20% HUVEC-conditioned media (HUVEC-CM) compared to mock control. The level of Wnt pathway activity induced with 20% HUVEC-CM was similar to that due to treatment with 2nM Wnt3a (N=11 batches of HUVEC-CM, each tested twice in triplicate). **(C)** HUVEC RT-PCR analysis revealed expression of Wnt 3, 7b, and R-spondin 3 (N=3). **(D)** In the context of fracture repair, callus macrophages (CD45+, F4/80+, CD11b+), endothelial cells (CD31+, Ve-Cadherin+), and chondrocytes (tdTomato+, CD45-) also expressed numerous Wnts (N=3/cell population). Although endothelial cells, followed by chondrocytes, expressed the greatest variety of Wnts, the CD45-cell population expressed the greatest quantity of Wnt ligands, as evidenced by **(E)** the relative expression of Wnt10b, the most highly expressed ligand (N=3/cell population). (*) = p< 0.05. (**) = p< 0.01.

In order to confirm that endothelial cells express Wnts in the biological context of fracture repair and to identify other potential sources of Wnts, gene expression analysis of Wnt ligands was performed on FACS-sorted callus macrophages, chondrocytes, and endothelial cells, as well as total callus CD45+ and CD45-cell populations. Endothelial cells (CD31+, Ve-Cadherin+), macrophages (CD45+, F4/80+, CD11b+), and total callus CD45+ and CD45-cell populations were isolated from C57BL/6 mice at 14 days post-fracture. Chondrocytes (tdTomato+, CD45-) were isolated from Aggrecan-Cre^ERT2^∷tdTomato HZE reporter mice at 10 days post-fracture in order to obtain as pure a population of chondrocytes as possible. The cartilage intermediate reaches its greatest size and the callus primarily consists of cartilage at 10 days post-fracture. All cell populations expressed numerous Wnt ligands, with endothelial cells followed by chondrocytes expressing the greatest number of Wnts and CD45+ cells expressing the fewest **(Fig 5, D-E).** Although endothelial cells expressed the greatest variety of Wnts, the total callus CD45-cell population expressed the greatest quantity of Wnts as demonstrated by the comparison of gene expression levels for Wnt 10b, the most highly-expressed Wnt within the fracture callus **(Fig 5, E)**.

### Fate of Callus Chondrocytes beyond Osteoblasts

Data from the loss- and gain-of-function experiments above suggest that activation of Wnt within chondrocytes serves as a molecular switch enabling these cells to become osteoblasts. However, fractures in lineage tracing experiments using tissue transplantation and Aggrecan-Cre^ERT2^-tdTomato mice suggest that not all chondrocytes become osteoblasts, but a portion also contribute cells to the lining of newly formed bone.^8,14^ These observations, coupled with our earlier observation that chondrocytes lacking β-catenin do not become bone **(Fig 1)**, but rather are sloughed into the marrow where they retain the ability to contribute to subsequent fracture healing **(Fig 2)**, we hypothesized that these chondrocytes may be highly plastic with multi-lineage potential.

We tested this secondary hypothesis by performing extended lineage tracing experiments using two approaches. First, we transplanted cartilage derived from the fracture callus of a Rosa26-(β-galactosidase-expression) mouse into a critical-sized defect as in the past,^14^ and second we performed a lineage analysis during fracture healing using the Aggrecan-Cre^ERT2^-tdTomato mouse.^8^ At day 35 post-transplantation the bone had healed and no cartilage was present in the callus in either experimental model. However, after transplantation we observed donor cells in newly formed cortical bone and in the periosteal and endosteal lining **(Fig. 6A)**. Similarly, at 35 days after fracture, genetic lineage tracing demonstrated tdTomato-positive cells in newly formed cortical bone and in the periosteal and endosteal lining of the bone **(Fig. 6E)**. To test whether the bone lining cells were capable of contributing to a new fracture callus, 35 days after transplantation or fracture, we created a second fracture in the same anatomical location. In both cases, cells within the original transplant and cells originally labeled with tdTomato contributed to the formation of new cartilage and bone within the callus at day 10 after injury **(Fig. 6, B, C, D, F, G, H)**. These data suggest that chondrocytes give rise to the periosteal and endosteal progenitor cells that heal injured bone.

**Figure 6:**
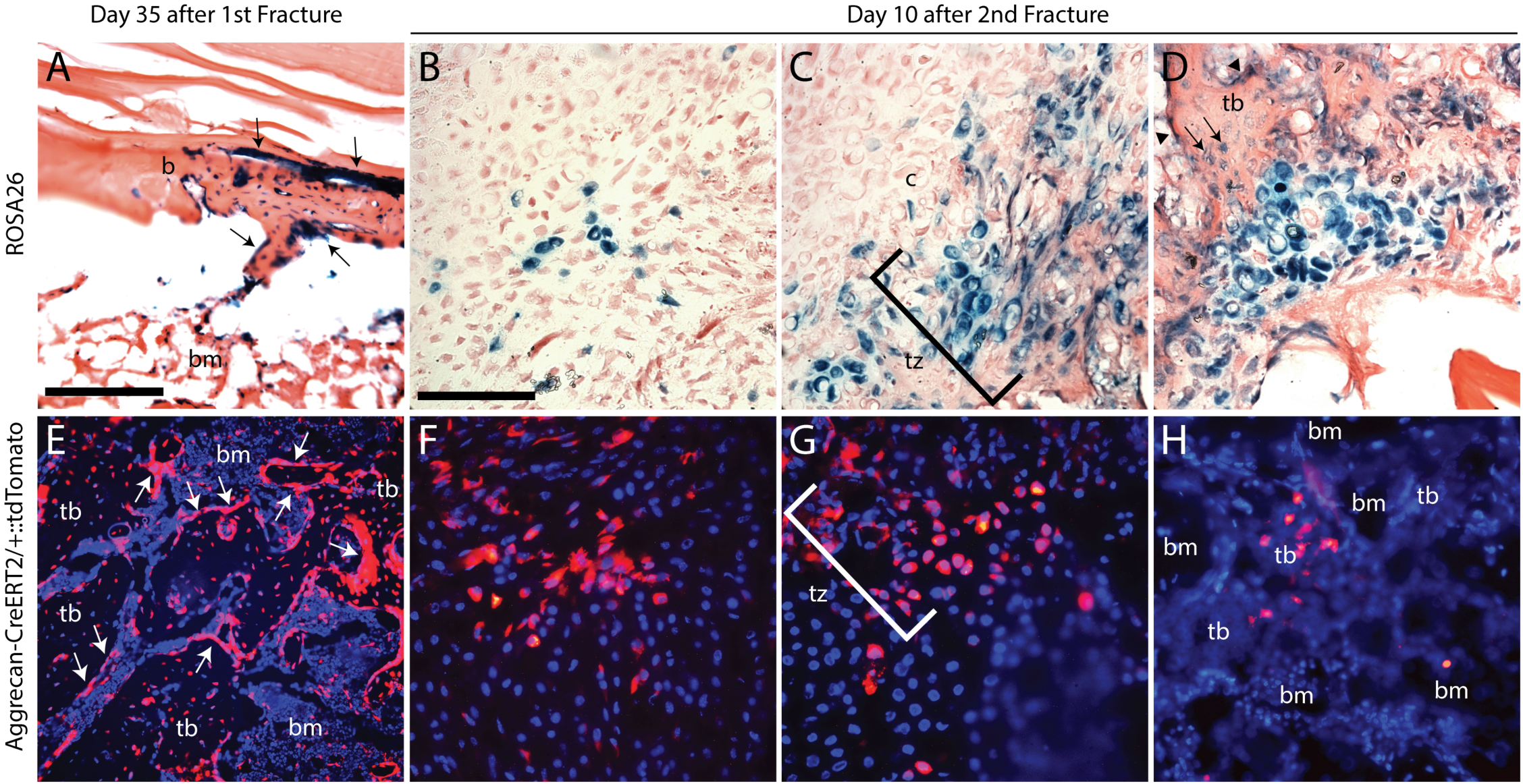
Lineage tracing during serial fracture. At 35 days after **(A)** transplantation of cartilage from ROSA26 mice (n=3) into a critical sized defect or fracture of Aggrecan-Cre^ERT2^-tdTomato mice (n=9), labeled cells are seen lining the surface of newly formed bone (arrows). After transplantation of ROSA26-derived cartilage and refracture (n=3), donor derived cells were found in **(C)** the newly formed cartilage, **(D)** chondrocytes and osteoblasts in the transition zone (tz), and **(D)** osteoblasts (arrows) in the newly formed trabecular bone (tb) and in cells lining newly formed bone (arrowheads). Similarly, after refracture of Aggrecan-Cre^ERT2^-tdTomato mice (n=9) tdTomato positive cells were observed in **(F)** cartilage, **(G)** cells comprising the transition zone (tz) and **(H)** cells in newly formed trabecular bone (tb). b, bone; tb, trabecular bone; bm, bone marrow, tz, transition zone. Scale bars: A,E, 200μm, B-D, F-H, 100 μm.

## Discussion

Numerous genetic lineage tracing studies have demonstrated that chondrocytes directly contribute to endochondral bone formation by transforming into osteoblasts.^8,12–17^ This has been shown in the contexts of bone development, postnatal growth, and fracture repair with an array of reporter systems (Col10-Cre, Col2-Cre^ERT2^, Aggrecan-Cre^ERT2^, eGFP- and β-galactosidase-labelled tissue transplants).^8,12–17^ We advanced this here and show that chondrocytes are also likely precursors of progenitor cells that line the newly formed bone. Our data also suggest that b-catenin signaling may act as a cell fate switch in the chondrocytes during fracture healing.

The canonical Wnt pathway has been repeatedly shown to promote osteogenesis in the context of intramembranous bone repair.^28–30^ Pathway inhibition through adenoviral expression of Dkk1 significantly reduces bone formation in transcortical defect models, which heal through intramembranous ossification.^28^ Conversely, activation of canonical Wnt signaling through deletion of pathway inhibitors (Sclerostin, Axin2) significantly increases intramembranous bone formation.^29^ However, the role of canonical Wnt signaling during endochondral healing has been less explored.

Interestingly, during development, the canonical Wnt pathway has been shown to act as a “molecular switch” by promoting osteogenic differentiation while repressing chondrogenic cell fate decisions.^31–33^ Inhibition of canonical Wnt signaling through deletion of β-catenin from skeletogenic mesenchyme results in early osteoblast differentiation arrest and ectopic cartilage formation.^31,33^ Similar results have been reported from *in vitro* experiments that inhibit canonical Wnt signaling in mesenchymal progenitor cells.^31^ Recently, Houben *et al.* demonstrated that canonical Wnt signaling also serves as a molecular regulator of chondrocyte-to-osteoblast transformation during endochondral bone development.^16^ Deletion of β-catenin from hypertrophic chondrocytes using a Col10a1-Cre resulted in significantly reduced bone formation, whereas induced expression of an indestructible form of β-catenin using the same Cre line produced osteopetrotic bone.^16^

Numerous Wnt ligands, receptors, and components of the transduction machinery have been shown to be expressed during endochondral healing, but their functional role remains unclear.^34,35^ Our data indicate that β-catenin and canonical Wnt signaling in fracture callus chondrocytes is critical to chondrocyte-to-osteoblast transformation during endochondral repair. Conditional deletion of β-catenin using an inducible Aggrecan-Cre resulted in significantly reduced new bone formation and increased cartilage retention. This confirms work from a similar study using Col2a1-ICAT transgenic mice to inhibit canonical Wnt signaling in chondrocytes that found delayed cartilage and reduced bone formation following tibia fracture.^36^ Importantly, our data indicate that inhibition of canonical Wnt signaling does not increase chondrocyte cell death. This confirms previously published evidence demonstrating that cell death is not the exclusive fate of hypertrophic chondrocytes.^8,16^ Indeed, our tdTomato lineage tracing analysis following deletion of β-catenin in callus chondrocytes revealed that tdTomato positive cells detach from their cartilage matrix and remain within the marrow cavity up to 35 days post-fracture and upon re-injury contribute to the cartilage callus.

It is possible that the prolonged presence of tdTomato+ cells within the marrow cavity is due to the phagocytic activity of hematopoietic cells that have engulfed chondrocytes. FACS analysis of tdTomato positive cells isolated from the marrow cavity following deletion of β-catenin revealed a subset of cells that were also positive for CD45, a marker of hematopoietic cell lineages.^25,26^ Significant dual tdTomato+ and CD45+ labelling was only observed upon deletion of β-catenin. Control marrow cells possessing the tdTomato reporter and wildtype copies of β-catenin had little to no dual-labelled cells. Furthermore, the number of dual-labelled cells found in control samples was similar to that found in wildtype mice that did not possess the tdTomato reporter. Thus, tdTomato and CD45 analysis were highly specific. Hematopoietic engulfment of detached chondrocytes is reminiscent of efferocytosis, a mechanism used by phagocytic cells to remove cellular debris for the resolution of tissue damage and during normal tissue development.^37,38^ Phagocytosis is mediated by both professional and non-professional phagocytic cells including macrophages, dendritic cells, tissue-infiltrating monocytes, neutrophils, and eosinophils as well as epithelial cells of mammary epithelium and astrocytes in the brain.^38–40^ However, in the majority of cases, phagocytic clearance is primarily undertaken by macrophages.^37^ This is likely the case in the context of hematopoietic engulfment of chondrocytes during fracture repair, since we show macrophages are present at the Transition Zone.

The canonical Wnt pathway is an important regulator of osteoclastogenesis.^41^ Inhibition of β-catenin in hypertrophic chondrocytes during bone development impairs trabecular bone formation and is associated with enhanced subchondral osteoclast number and increased expression of *receptor activator of nuclear factor kappa-B ligand* (*Rankl*), which is known to stimulate osteoclastogenesis.^41^ Houben *et al.* demonstrated that modulation of canonical Wnt signaling in chondrocytes affects osteoclastogenesis by altering the expression of *Rankl* and *Osteoprotegrin* (*Opg*), a Rankl antagonist.^16^ They also demonstrated that downregulation or increased activation of osteoclastogenesis was able to partially rescue the phenotypic changes observed from loss or stabilization of β-catenin, respectively.^16^ Our data indicate that β-catenin and the canonical Wnt pathway may play a similar role during endochondral repair. TRAP staining revealed that deletion of β-catenin increased the localization of TRAP-positive cells to the border of the cartilage callus. This localized increase in osteoclastic activity may be responsible for the robust detachment of tdTomato+ cells observed within the marrow cavity.

In contrast to deletion of β-catenin signaling in callus chondrocytes, pathway gain-of-function through conditional expression of an indestructible form of β-catenin resulted in increased and accelerated bone formation. This increase in bone formation corresponded with a simultaneous decrease in callus cartilage composition, indicating that canonical Wnt signaling is responsible for regulating chondrocyte-to-osteoblast transformation. Chondrocyte transformation is known to occur in a specific region of the fracture callus, which we have termed the “Transition Zone” as this is where the callus cartilage joins the newly formed bone.^8,14^ Previously we showed that hypertrophic chondrocytes at the Transition Zone express transcription factors traditionally associated with stem cell pluripotency (Sox2, Oct4, Nanog).^8^ Importantly, we demonstrated that *Sox2* plays a functional role in fracture repair with deletion of *Sox2* during the cartilaginous phase of healing resulting in significantly reduced new bone formation and increased cartilage retention within the fracture callus.^8^ It is unlikely that chondrocytes truly become pluripotent during transformation, but we suggest that *Sox2* may play a role in chondrocyte plasticity. Recently, Willet *et al.* coined the term “paligenosis” to describe the process by which differentiated cells revert to a proliferative and regenerative state and undergo metaplasia in response to tissue injury in the context of stomach and pancreatic regeneration.^42^ Our data suggest a similar process occurs in the context of fracture repair.

Our data support the concept that vascular invasion is a critical regulator of metaplasia during endochondral fracture repair. Secreted factors from human umbilical vein endothelial cells (HUVECs) induce expression of pluripotency transcription factors (*sox2, oct4, nanog*) and the osteogenic gene o*steocalcin* in fracture callus explants.^8^ Here we demonstrated that HUVEC-secreted factors are also capable of stimulating canonical Wnt signaling using the luciferase-based 293-STF Wnt reporter cells, and qPCR analysis of HUVECs reveals that these cells express numerous Wnts and R-Spondins. Cells isolated from fracture calli by FACS also revealed significant expression of a variety of Wnt ligands. Endothelial cells, macrophages, and chondrocytes were isolated along with total callus CD45+ and CD45-cell populations. All populations expressed numerous Wnt ligands, with CD45-cells expressing a significantly larger quantity and variety of Wnts compared to CD45+ cells and with endothelial cells followed by chondrocytes expressing the most Wnts out of the three specifically-isolated cell types. Taken together these data suggest that multiple cell types may contribute to the production of Wnt ligands for promoting the osteogenic switch within chondrocytes.

Together, our data demonstrate the critical role of β-catenin/Wnt signaling to the process of chondrocyte-to-osteoblast transformation and the contribution of callus chondrocytes to not only new bone formation but also to the progenitor cell populations found within the periosteum and endosteum. These data provide important insight towards a complete understanding of the process of endochondral repair. **Figure 7** outlines our current understanding of this process to date.

**Figure 7:**
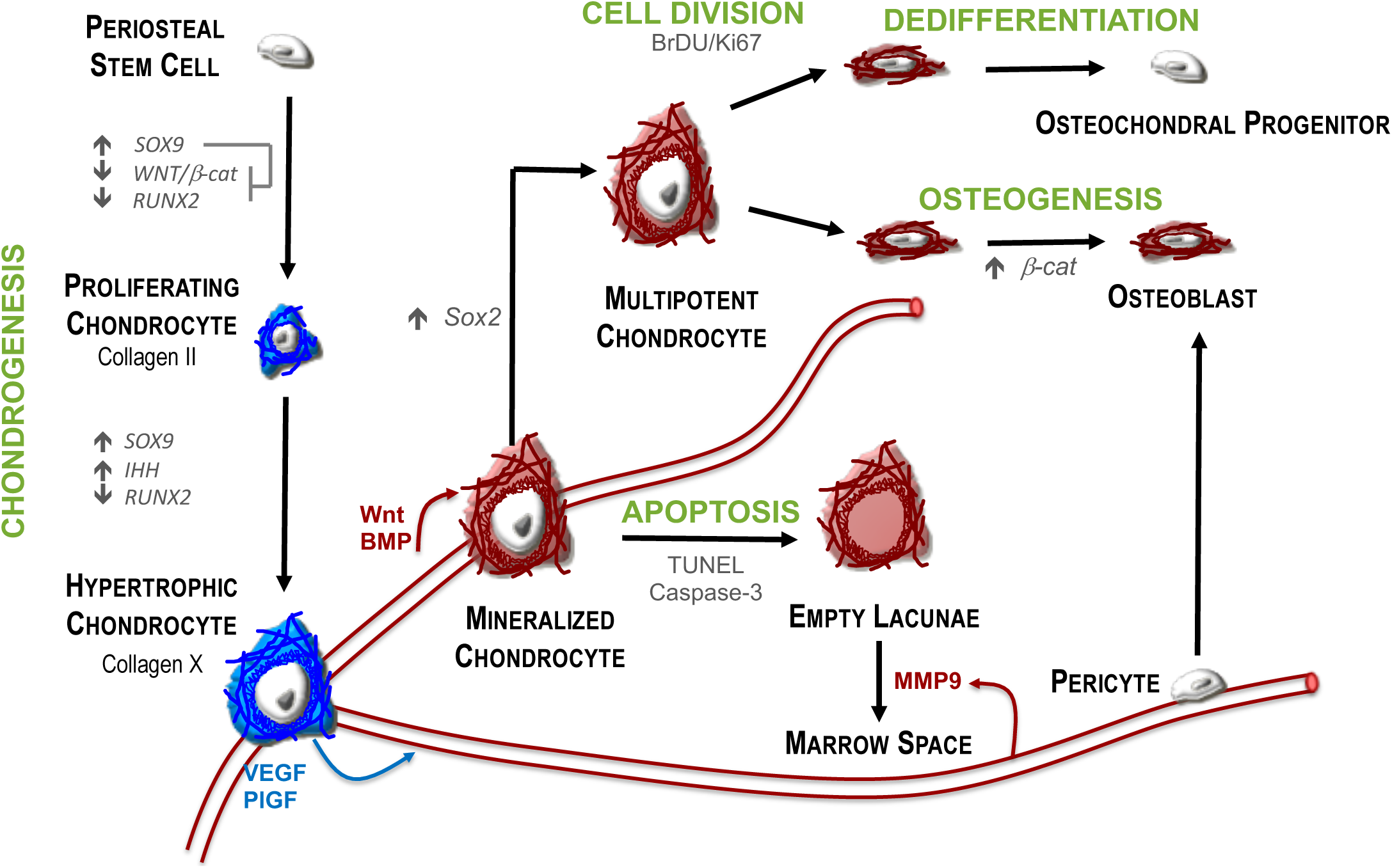
Cell fate decisions during endochondral repair. Evidence from this study contributes to our greater understanding of endochondral repair. This process begins with the differentiation of periosteal osteochondral stem cells, which differentiate into chondrocytes that proliferate to form the cartilage callus. Chondrocytes then undergo hypertrophy, recruit the vasculature and mineralize their surrounding matrix. A significant population of hypertrophic chondrocytes survive to gain functional cellular plasticity through the expression of *Sox2*. The osteogenic fate of these cells is determined through activation of β-catenin signaling. In the absence of this osteogenic switch, a portion of chondrocytes undergo apoptosis in order to make room for the marrow cavity, while another sub-population contribute to the periosteum and endosteum of the newly formed bone with the capacity to contribute to future fracture repair.

## Conclusion

Our data demonstrate the key role of β-catenin and the canonical Wnt pathway in chondrocyte-to-osteoblast transformation during endochondral repair. We show that canonical Wnt signaling is not required for chondrocyte survival but rather that this pathway likely functions to regulate gene expression for osteogenic programing and osteoclastogenesis. Furthermore, we present evidence not only confirming the contribution of chondrocytes to new bone formation, but also demonstrating that they contribute to the progenitor cell populations found within the periosteum and endosteum. Together this data significantly advances our understanding of endochondral repair and provides novel insight into the development of Wnt-based fracture therapies.

## Materials and Methods

### Fractures

All studies were approved by the UCSF Institutional Animal Care and Use Committee. The following mice were obtained from Jackson Labs: Aggrecan-Cre^ERT2^ (Stock #: 019148), β-Catenin floxed (Stock #: 004152), tdTomato HZE (Stock #: 007909), C57BL/6J (Stock #: 000664), and ROSA26 (Stock#:). Immunocompromised SCID mice were obtained from Charles River. Cttnb1GOF^43^ mice were obtained from our collaborators at UCSF and maintained in our colony for the duration of this study. Genotype was confirmed for all mice through gel electrophoresis according to Jackson Labs and donor lab’s genotyping protocols. Adult (10-18 week) male mice were anesthetized with a 1:1 mixture of ketamine (60 mg/kg) and dexmedetomidine (0.3 mg/kg). Closed non-stable fractures were created in the mid-diaphysis of the right tibia through 3-point bending.^44^ Fractures were not stabilized to promote robust endochondral repair. Animals were revived using an atipamezole reversal agent (6 mg/kg) and were given buprenorphine analgesic (0.05-0.1 mg/kg) immediately post-op, at 4 and 24 hours post-fracture, and as needed. Cre recombination was induced with daily injections of Tamoxifen (75 mg/kg in corn oil, Sigma, #T5648-5G) from days 6 to 10 post-fracture.^8^ Mice were monitored and allowed to ambulate freely until harvest at 7, 10, 14, and 21 days post-fracture (N>5/group/time point). To identify any potential differences in fracture repair based on sex, female mice were also harvested at 14 days post-fracture (N>5/group).

### Histology

Fractured tibiae were fixed in 4% paraformaldehyde (PFA, pH 7.4) for up to 24 hours at 4C and decalcified for 2 weeks in 19% ethylenediaminetetraacetic acid (EDTA, pH 7.4). EDTA was replenished 5 times per week. Tibiae were either paraffin or cryo-embedded. Tissues were sectioned (10 µm) for histological and fluorescence analysis. Halls Brunt Quadruple (HBQ) histology was used to visualize cartilage and bone (cartilage stains blue, bone stains red). N>5/group/time point.

### Stereology

Quantification of callus size and composition (cartilage, bone, fibrous, marrow space) was determined using an Olympus CAST system (Center Valley, PA) and software by Visiopharm (Hørsholm, Denmark) according to established methodologies.^45^ HBQ stained serial sections through the entire leg, spaced 300 µm apart were analyzed. The fracture callus was outlined using low magnification (20x, 2X objective with 10X ocular magnification) to determine the region of interest. 15% of this region was quantified using automated uniform random sampling to meet or exceed the basic principles of stereology.^45^ Cell and tissue identity within each random sampling domain was determined at high magnification (200X, 20X objective with 10X ocular magnification) according to histological staining patterns and cell morphology. Absolute volumes of specific tissues (e.g. bone or cartilage) were determined using the Cavalieri formula and these absolute volumes were used to determine the relative percent composition of tissues within the fracture callus.^45^ Total fracture callus volume was measured as callus points. Marrow space was identified as tissue that fell within a blood vessel or marrow cavity of the new bone.

### TUNEL Assay

The Roche In Situ Cell Death Detection Kit (Roach, #11687959) was used according to the manufacturer’s protocol. Sections were deparaffinized, treated with proteinase K (20 ug/ml in 10 mM Tris-HCl, pH 8, 15 min), and then reacted with the kit for 1 hour at 37C in the dark. Positive controls were treated with DNase I prior to TUNEL reaction, while negative controls were not treated with the TUNEL reaction enzyme. Slides were mounted in VectaShield with DAPI (Vector, #H-1200) and visualized using an epifluorescence microscope (Leica DM500B).

### Lineage Tracing

Aggrecan-Cre^ERT2/+^∷β-catenin^fl/fl^∷tdTomatoHZE^/+^ mice were used to assess chondrocytes following deletion of β-catenin. Unstable fractures were created and tamoxifen was administered as above. Samples were harvested 14 and 35 days post-fracture, fixed and decalcified as above, cryoembedded, and sectioned (10 µm). Samples were mounted in VectaShield with DAPI (Vector, #H-1200) and visualized using an epifluorescence microscope. N=5/time point. To assess the contribution of callus chondrocytes to the development of future fracture calli, a separate set of mice received a second injury 35 days after the first fracture and the contribution of tdTomato cells was assessed 10 days later (N=9).

Similar lineage tracing analysis was performed in control animals (Aggrecan-Cre^ERT2/+^∷tdTomatoHZE^/+^). Tamoxifen was administered as above. Tissues were examined 35 days after fracture via fluorescent microscopy (n=9). A separate set of animals received a second fracture 35 days after the initial injury, and the presence and distribution of tdTomato positive cells was assessed 10 days later (n=9).

### Cartilage Transplantation and Re-Fracture

Cartilage grafts were isolated from the central portion of day 7 fracture calli harvested from ROSA26 reporter mice *ex vivo* by using a microscope to dissect out the cartilage and remove all non-cartilaginous adherent tissues and the perichondrium as previously described.^14^ Grafts were transplanted into 2 mm critical-sized defects created by osteotomy in SCID mice immediately after dissection. An 8.0 suture was used to secure graft in place by closing the muscle. Tibiae were externally stabilized with a customized circular fixator consisting of two 2 cm circular rings held concentrically by three threaded rods. This device provides rigid fixation and has been extensively described previously.^14,46^ Animals were harvested at 35 days post-fracture (n=3), other animals (n=3) received a second fracture within the original fracture site was made by 3-point bending as above^44^ at this time, and ROSA26 cells were analyzed in 10 days later.

### Immunohistochemistry

Immunohistochemistry for F4/80 (eBiosciences, #144801, 1:100) was performed on paraffin-embedded sections of tibiae harvested 10 days post-fracture (N>3). The basic protocol included endogenous peroxidase blocking in 3% H_2_O_2_ (30 min, RT), antigen retrieval in 0.1% Trypsin (20 min sat 37C then 20 min at RT), and non-specific epitope blocking with 5% goat serum (1 hr, RT). Primary antibodies were applied to sections overnight at 4C. Biotinylated species-specific secondary antibodies were diluted in PBS with 5% GS and applied to tissues for 1 hr at room temperature (Santa Cruz, #sc-2041, 1:250). Staining was detected using the VectaStain ABC Elite Kit (#PK-6100) and 3,3’-diaminobenzidine (DAB) colorimetric reaction.

### TRAP Staining

TRAP staining was performed on paraffin sections using the Sigma TRAP staining kit (#387A) according to the manufacturer’s protocol. Slides were mounted with Permount and visualized with brightfield or DIC microscopy.

### HUVEC Conditioned Media

Human umbilical vein endothelial cells (HUVECs) were cultured in Medium 200 (Gibco, #M-200-500) with Large Vessel Endothelial Supplement (LVES 50X, GIBCO, #A14608-01), 1% Penicillin/Streptomycin, and 10% MSC FBS (Gibco, #12662-029) until ∼70% confluent. Cells were washed twice with dPBS and cultured for an additional 48 hours without FBS. Conditioned media was filtered (0.22µm) and stored at −80C. HUVECs were harvested into TRIzol (Invitrogen, #15596026) and stored at −80C. N=11 media harvests.

### Wnt Reporter Assay

293-STF luciferase Wnt reporter cells were obtained from the Christopher Garcia laboratory.^47^ Cells were seeded at 20,000 cells/well in a 96-well plate and cultured overnight in DMEM (Gibco, #10566-016), 10% FBS, 1% Pen/Strep. Cells were treated with 20% HUVEC-conditioned media (HUVEC-CM), 2 nM Wnt3a, or with unconditioned media for 24 hours. All samples were run in technical triplicates and received 25 nM Rspon2-Fc. The Dual-Luciferase Reporter Assay System (Promega, #E1960) was used to assess luciferase activity.

### RT-PCR

Fracture calli were harvested 10 days post-fracture from β-catenin gain-of-function and control mice in order to assess Axin2 gene expression and confirm over-activation of the canonical Wnt pathway. Callus tissues were isolated from surrounding muscle and cortical bone and homogenized in TRIzol (Invitrogen, #15596026). RNA isolation was performed according to the manufacturer’s protocol followed by DNase treatment (Invitrogen, #Am1906). cDNA was reverse transcribed with iScript cDNA Synthesis Kit (Bio-Rad, #1708890) and quantitative RT-PCR was performed using SYBR-based primers (**Supplemental Table 1)** and RT SYBR Green qPCR Mastermix (Qiagen, #330509) on a CFX96 Touch Thermal Cycler (Bio-Rad) run to 50 cycles (N=3-5/treatment group). RNA isolation, DNase treatment, and reverse transcription was performed as above for Wnt expression analysis on HUVEC cells. However, TaqMan probes **(Supplemental Table 2)** and TaqMan Universal Master Mix II with UNG (#4440038) were used. Analysis was run on a CFX96 Touch Thermal Cycler (Bio-Rad) to 45 cycles (N=3). All samples were run in triplicate and relative gene expression was calculated by normalizing to the housekeeping gene GAPDH (2^-**Δ**CT^). Fold change was calculated as 2^-**ΔΔ**CT^.

### FACS

Macrophages (CD45+, F4/80+, CD11b+), endothelial cells (CD31+, Ve-Cadherin+), and total callus CD45+ and CD45-cell populations were isolated from C57BL/6 mice at 14 days post-fracture, whereas chondrocytes (tdTomato+, CD45-) were isolated from Aggrecan-CreERT2^/+^∷tdTomato HZE^/+^ reporter mice at post-fracture day 10 after tamoxifen injection from days 6-10. A defined area around the fracture callus was isolated, weighed, skin and muscle eliminated and bone marrow discarded or stored separately. The remainder of the fracture callus was minced and disassociated manually in Collagenase/Dispase (1 µg/ml, Roche, #10269638001) followed by digestion for 1 hour at 37 degrees. After digestion the cells were passed through a 100µm filter and rinsed with PBS. Cells were then pelleted and counted with a hemocytometer. Isolated cells were blocked for 10 minutes at room temperature with anti-FcR antibodies (ATCC, 24G2 hybridoma) and then stained directly with conjugated fluorescent antibodies **(Supplemental Table 3)**. Red Dead or Ghost Violet (Biosciences, #13-0863-T100) was used for the detection of dead cells. Isotype controls and fluorescence minus one controls were used to gate for background staining. Cells were sorted on a FACSAriaIII (BD Biosciences, San Jose, CA) and FlowJo Software 9.6 (Treestar, Ashland, OR) was used for analysis. N=3/population.

### Fluidigm Gene Expression Analysis

Total RNA was isolated from FACS-sorted cells using the Arcturus PicoPure RNA Isolation Kit (Thermo, #12204-01) and was DNase treated (Ambion DNA-Free: DNase Treatment & Removal Kit, #AM1906) according to the manufacturer’s protocols. RNA was reverse transcribed into cDNA using Maxima H Minus Reverse Transcriptase (Thermo, #EP0752) followed by RNase treatment with RNAse H (Thermo, #18021071). Pre-amplification of cDNA was performed using the Fluidigm Pre-Amp MasterMix (Fluidigm, #100) and TaqMan assays **(Supplemental Table 2)** according to manufacturer’s instructions. qPCR for GAPDH and β-actin was performed in technical triplicates for all samples using a Viia7 thermocycler (Invitrogen) to determine optimal cDNA sample concentration. Complete gene expression analysis was performed for all samples using a BioMark 48.48 dynamic array nanofluidic chip (Fluidigm Inc., BMK-M-48.48, #68000088). Ct values were normalized to the housekeeping gene GAPDH. N=3/cell population.

### Statistics

Data plots were generated using GraphPad Prism 8. Initial analysis of variance was assessed using RStudio and statistical significance was determined with Prism. The non-parametric Mann-Whitney-Wilcoxon test was used when comparing two groups. ANOVA followed by post-hoc multiple-comparisons analysis through Tukey’s HSD test was used when comparing more than two groups. P≤0.05 was designated as statistically significant.

## Acknowledgements

We would like to acknowledge Charles Lam for his assistance with mouse breeding and genotyping, Alyson Fiorentino for her assistance with statistical analysis, and Drs. Chris Garcia and Claudia Janda who generously provided the reporter cells and equipment to perform the Wnt reporter assay for this study. WntGOF mice expressing an indestructible form of β-catenin were gifted by Dr. Ophir Klein. We would also like to thank Gina Baldoza and Anna Lissa Wi at the UCSF Orthopaedic Trauma Institute for their service in daily lab function and grants administration.

## Author contributions

SAW, DH, TM, RSM, and CSB devised the experiments. SAW, DH, TS, EN, and OB executed the experiments. EB, BM, DH, ECH, and MN provided expert advice in executing experiments. SAW performed data analysis. SAW, TM, ECH, MN, CSB, and RSM provided financial support and materials for the studies. SAW drafted and all authors contributed to editing the manuscript.

---

**Figure S1:**
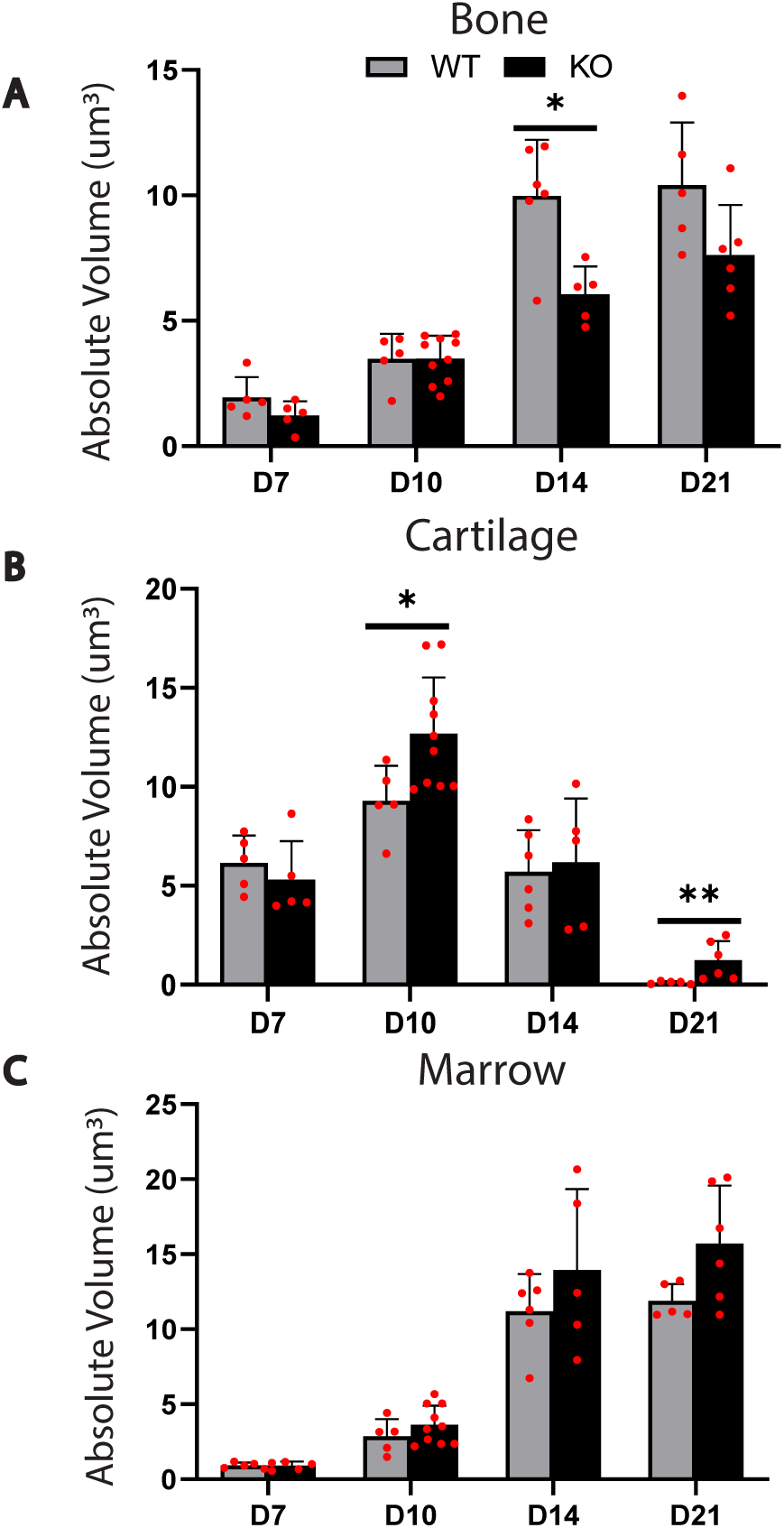
Stereological Quantification of Callus Tissue Components Following Deletion of β-catenin in Chondrocytes. **(A-C)** Stereological quantification of HBQ histology reveals that inhibition of canonical Wnt signaling through conditional deletion of β-catenin in chondrocytes significantly reduces **(A)** absolute volume of trabecular bone and increases **(B)** absolute volume of cartilage retention. There is a trend in **(C)** increasing marrow absolute volume with deletion of β-catenin. However, this difference is not statistically significant. N=6-10/group/time point. (*) = p< 0.05. (**) = p< 0.01.

**Figure S2:**
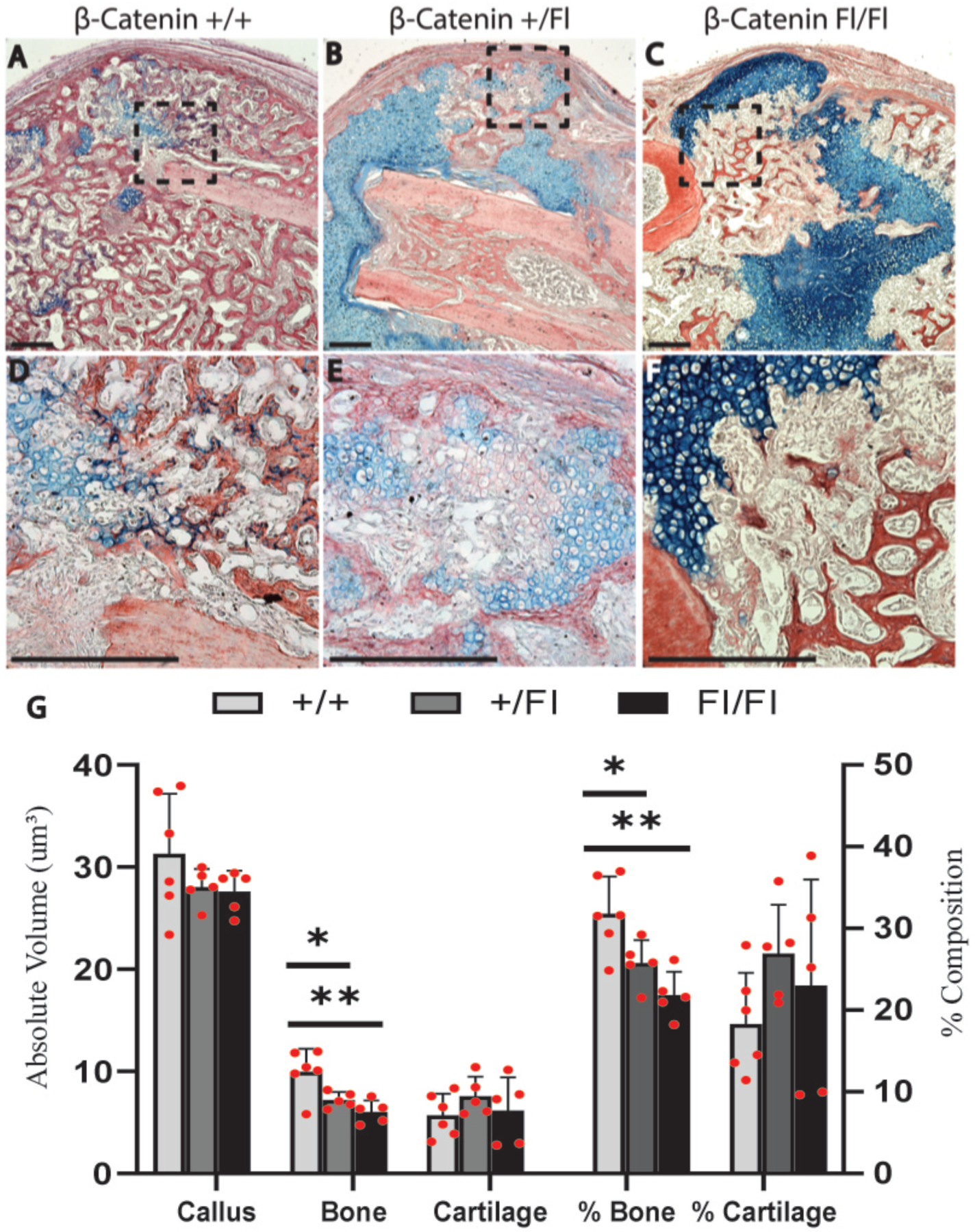
Statistically Significant Reduction in Bone Formation Results from Deletion of Either One or Both Copies of β-catenin in Chondrocytes. **(G)** Stereological quantification of **(A-F)** HBQ histology reveals that inhibition of canonical Wnt signaling through conditional deletion of either **(B**,**E)** one or **(C**,**F)** both copies of β-catenin in chondrocytes results in a statistically significant reduction in absolute bone volume and % bone composition. A greater reduction is observed with the loss of both copies. **(D-F)** High magnification of the boxed regions in **(A-C)** depicts changes in Transition Zone morphology. HBQ histology: cartilage = blue, bone = red. N=5-6/group/time point. Scale = 1000μm. (*) = p< 0.05. (**) = p< 0.01.

**Figure S3:**
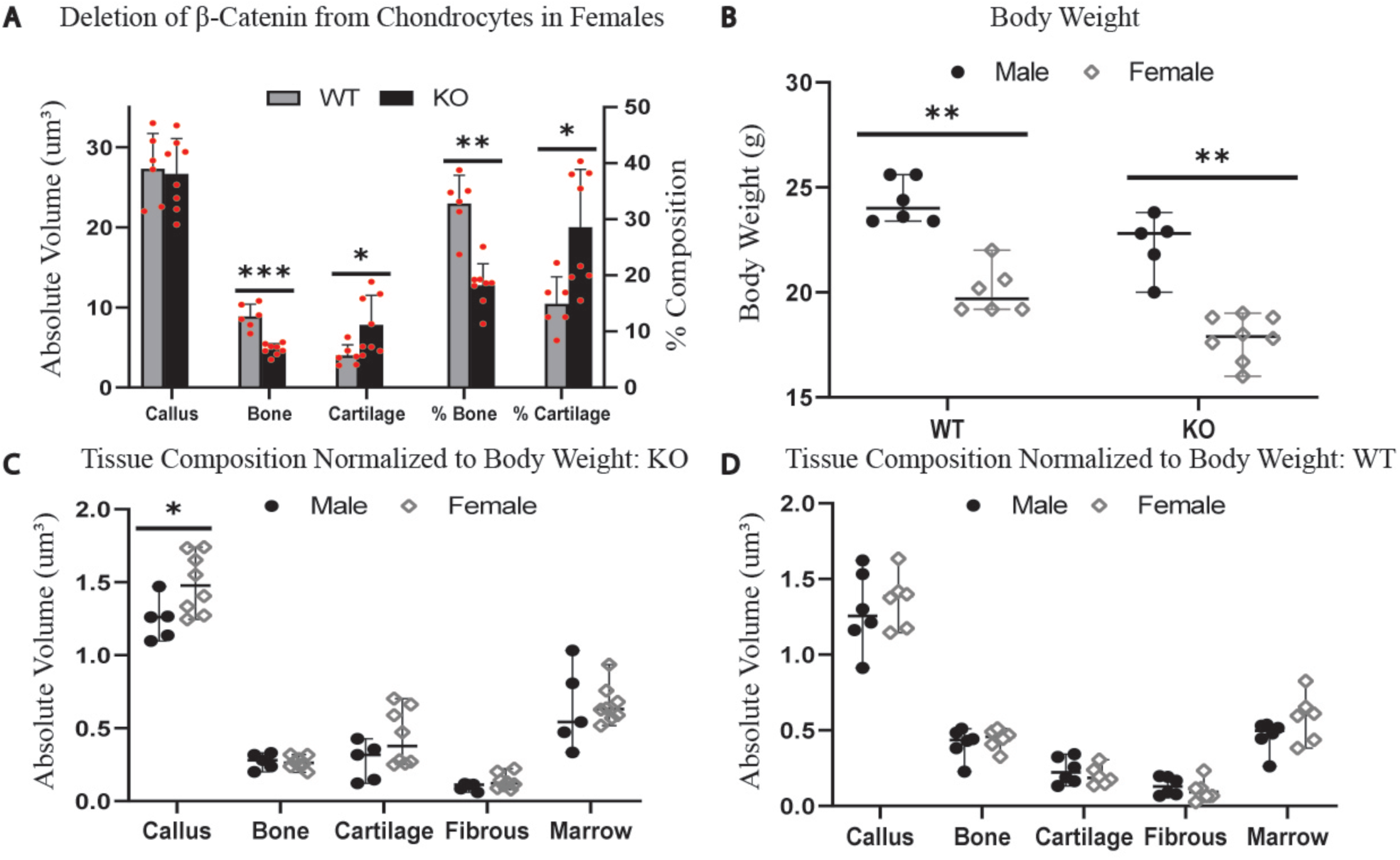
Canonical Wnt Signaling Regulates Chondrocyte-to-Osteoblast Transformation in Both Males and Females. **(A)** Stereological analysis of fractures harvested from female mice at 14 days post-fracture reveals that inhibition of canonical Wnt signaling in chondrocytes through deletion of β-catenin significantly reduces bone formation and increases cartilage retention compared to controls. **(B)** Although female mice have a significantly lower body weight compared to male mice, there is **(D)** no statistically significant difference in tissue composition between sexes for control mice when normalized to body weight. **(C)** There is a slight increase in total callus size in female mice compared to males when β-catenin is deleted. However, all other tissue parameters show no statistically significant difference. N=6-8/group/time point. (*) = p< 0.05. (***) = p< 0.001.

**Figure S4:**
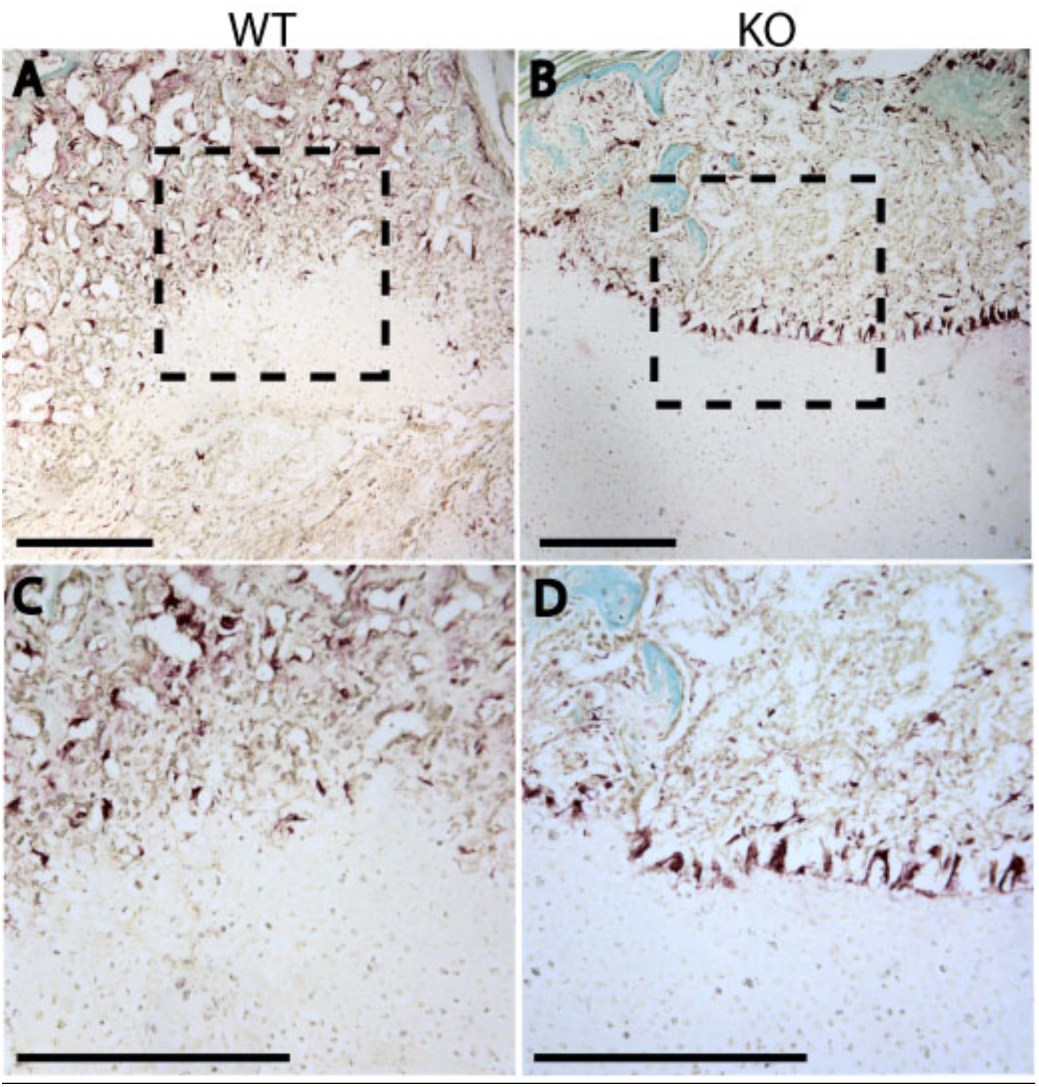
TRAP Staining Reveals Increased Localization of Osteoclasts to the border of the Cartilage Callus after Deletion of β-catenin in Chondrocytes. **(A-D)** TRAP staining reveals increased localization of osteoclasts at the cartilage callus border following **(B**,**D)** deletion of β-catenin in chondrocytes compared to **(A**,**C)** controls. **(C-D)** High magnification of the boxed regions in **(A-B)** depicts changes in Transition Zone osteoclast localization. Fast Green counter stain. N=5. Scale = 200μm.

**Figure S5:**
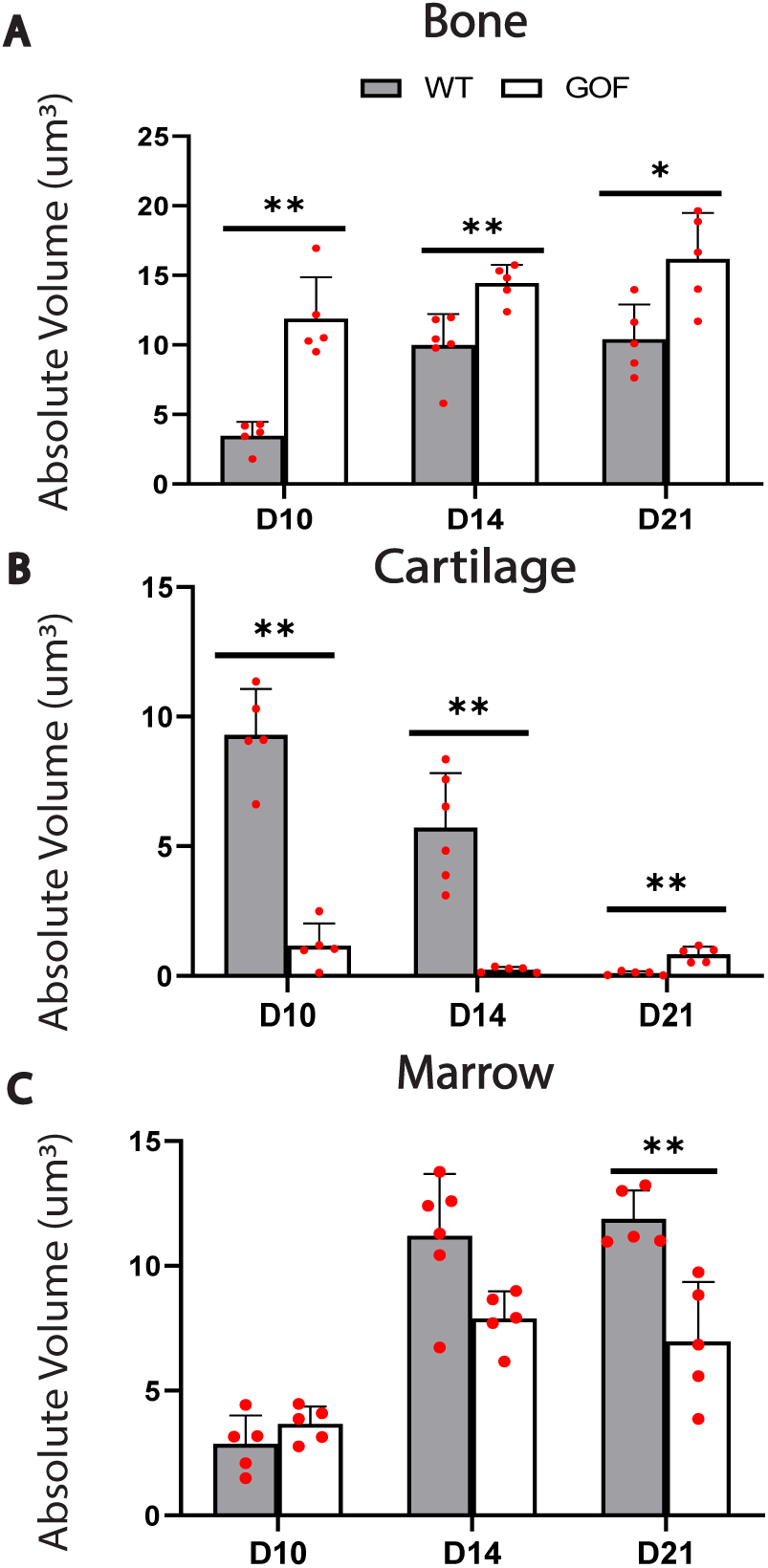
Stereological Quantification of Callus Tissue Components Following Over-Activation of Canonical Wnt Signaling in Chondrocytes. **(A-C)** Stereological quantification of HBQ histology reveals that over-activation of canonical Wnt signaling through conditional stabilization of β-catenin in chondrocytes **(A)** significantly accelerates and increases absolute trabecular bone volume. This increase in bone volume occurs simultaneously with **(B)** a reduction in absolute cartilage volume and **(C)** an increase in absolute marrow volume. N=6/group/time point. (*) = p< 0.05. (**) = p< 0.01.

