## supplemental data for "β-catenin Signaling Regulates Cell Fate Decisions at the Transition Zone of the Chondro-Osseous Junction During Fracture Healing"

**Supplementary Information:**

**Supplemental Table 1: SYBR Green Primers for Gene Expression Analysis**

| Primer | Species | Company | Sequence |
|---|---|---|---|
| Axin2 Forward | Mouse | Thermo | 5'- CTCCCCACCTTGAATGAAGA |
| Axin2 Reverse | Mouse | Thermo | 5'- ACTGGGTCGCTTCTCTTGAA - |

**Supplemental Table 2: TaqMan Assays Used for Gene Expression Analysis**

| Gene | Species | Company | Catalog # | Assay ID |
|---|---|---|---|---|
| 18S | Human | Thermo | 4331182 | Hs99999901_s |
| beta-actin (ACTB) | Mouse | Thermo | 4453320 | Mm01205647_ |
| GAPDH | Human | Thermo | 4453320 | Hs02758991_g |
| GAPDH | Mouse | Thermo | 4331182 | Mm99999915_ |
| Rspo1 | Mouse/Human | Thermo | 4453320 | Mm00507077_ |
| Rspo2 | Mouse | Thermo | 4453320 | Mm00555790_ |
| Rspo2 | Human | Thermo | 4448892 | Hs04400416_m |
| Rspo3 | Mouse/Human | Thermo | 4453320 | Mm01188251_ |
| Rspo4 | Mouse/Human | Thermo | 4453320 | Mm00615419_ |
| Wnt1 | Mouse/Human | Thermo | 4453320 | Mm01300555_ |
| Wnt10a | Mouse/Human | Thermo | 4331182 | Mm00437325_ |
| Wnt10b | Mouse/Human | Thermo | 4453320 | Mm00442104_ |
| Wnt11 | Mouse/Human | Thermo | 4453320 | Mm00437328_ |
| Wnt16 | Mouse | Thermo | 4453320 | Mm00446420_ |
| Wnt16 | Human | Thermo | 4453320 | Hs00365138_m |
| Wnt2 | Mouse/Human | Thermo | 4453320 | Mm00470018_ |
| Wnt2b (Wnt13) | Mouse/Human | Thermo | 4453320 | Mm00437330_ |
| Wnt3 | Mouse | Thermo | 4453320 | Mm00437336_ |

| | | | | |
|---|---|---|---|---|
| Wnt3 | Human | Thermo | 4448892 | Hs00902257_r |
| Wnt3a | Mouse | Thermo | 4453320 | Mm00437337_ |
| Wnt3a | Human | Thermo | 4453320 | Hs00263977_r |
| Wnt4 | Mouse/Human | Thermo | 4453320 | Mm01194003_ |
| Wnt5a | Mouse/Human | Thermo | 4453320 | Mm00437347_ |
| Wnt5b | Mouse/Human | Thermo | 4453320 | Mm01183986_ |
| Wnt6 | Mouse/Human | Thermo | 4448892 | Mm00437351_ |
| Wnt7a | Mouse/Human | Thermo | 4453320 | Mm00437356_ |
| Wnt7b | Mouse | Thermo | 4453320 | Mm01301717_ |
| Wnt7b | Human | Thermo | 4453320 | Hs00536497_r |
| Wnt8a | Mouse/Human | Thermo | 4453320 | Mm01157914_ |
| Wnt8b | Mouse/Human | Thermo | 4453320 | Mm00442108_ |
| Wnt9a | Mouse/Human | Thermo | 4453320 | Mm00460518_ |
| Wnt9b | Mouse/Human | Thermo | 4453320 | Mm00457102_ |

**Supplemental Table 3: Antibodies Used for FACS**

| **Target** | **Fluorofore** | **Company & Catalog #** |
|---|---|---|
| CD11b | PeCy7 | BD #552850 |
| CD31 | FITC | Biolegend #102406 |
| CD45 | FITC | BD #553080 |
| F4/80 | APC | eBioscience #17480182 |
| Ve-Cadherin (CD144) | PeCy7 | Biolegend #138016 |
| VEGFR2 (CD309) | APC | Biolegend #136405 |
